# Alpha and beta rhythms differentially support the effect of symbols on visual object recognition

**DOI:** 10.1101/2021.06.07.447387

**Authors:** Piermatteo Morucci, Francesco Giannelli, Craig Richter, Nicola Molinaro

## Abstract

Hearing spoken words can enhance visual object recognition, detection and discrimination. Yet, the mechanisms that underpin this facilitation are incompletely understood. On one account, words do not bias early visual processing, but rather affect later semantic or decision-making stages. However, recent proposals suggest that words can alter early visual processes by activating category-specific priors in sensory regions. A prediction of this account is that top-down priors evoke changes in occipital areas in anticipation of visual stimuli. Here, we tested the hypothesis that neural oscillations serve as a mechanism to activate language-generated visual priors. Participants performed a cue-picture matching task where cues were either spoken words, in their native or second language, or natural sounds, while their EEG and reaction times were recorded. Behaviorally, we replicated the previously reported label-advantage effect, with images cued by words being recognized faster than those cued by natural sounds. A time-frequency analysis of cue-target intervals revealed that this behavioral label-advantage was associated with enhanced power in posterior alpha (9-11 Hz) and beta oscillations (17-19 Hz), both of which were larger when the image was preceded by a word compared to a natural sound. Importantly, object recognition performance improved with high alpha power but slowed down with enhancement of beta synchronization. These results suggest that alpha and beta rhythms play distinct functional roles to support language-mediated visual object recognition: alpha might function to amplify sensory priors in posterior regions, while beta may (re)activate the network states elicited by the auditory cue.

## 1. Introduction

Hearing certain natural sounds (e.g., the croak of a frog) appears to automatically activate conceptual knowledge, enabling the perceptual system to quickly identify objects in the surroundings (e.g., the presence of a frog). Learning such cross-modal associations represents a crucial prerequisite for mediating interactions with the environment. In humans, conceptual representations can also be activated via language (e.g., “frog”). However, unlike natural sounds, linguistic symbols are categorical, making them uniquely suited to activate semantic information in a format that transcends within-category differences. It remains unclear whether phylogenetically young systems like language exert effects on perception similar to natural sounds, and what brain dynamics might support such effects. In the present study, we test the hypothesis that language boosts visual processes by sharpening categorical priors via the modulation of alpha/beta oscillations.

Conceptual representations activated by auditory cues have been shown to interact with the visual system in different ways. For instance, hearing words and natural sounds can rapidly drive visual attention towards specific entities in a scene (Huettig and Altmann, 2007); facilitate the recognition and discrimination of congruent object categories (Edmiston and Lupyan, 2015; Boutonnet and Lupyan, 2015); lower the detection threshold for ambiguous objects (Lupyan and Ward, 2013; Ostarek and Huettig, 2017); and even cause sensory illusions (Dils and Boroditsky, 2010). While this body of evidence suggests that both linguistic and non-linguistic cues activate content-specific representations, it is less clear whether these cues activate the *same* representations. Studies directly targeting this issue have often reported a “label-advantage” effect, that is, a facilitation when object recognition is preceded by words compared to non-linguistic cues (Lupyan and Thompson-Schill, 2012; Edmiston and Lupyan, 2015). This result suggests that language provides a particularly powerful tool to enhance visual processing.

To achieve these facilitatory effects on visual perception, linguistic categories could theoretically follow two possible pathways (Simanova et al., 2016). Language might not bias perceptual processes at early levels, but rather interact at later semantic or categorical decision-making stages (Gleitman and Papafragou, 2005; Klemfuss et al., 2012; Firestone and Scholl, 2014). On an alternative account, words could affect visual processing by setting categorical priors that alter early perceptual processing (e.g., Thierry et al., 2009; Kok et a., 2014; Boutonnet and Lupyan, 2015). Support for the latter account comes primarily from EEG studies showing that better recognition of images preceded by congruent words was associated with modulations of early event related potentials (ERPs) such as the P1 (Boutonnet and Lupyan, 2015, Noorman et al., 2018) – putatively considered an electrophysiological index of low-level visual processes (Spehlmann, 1965). Yet, these ERP experiments targeted the perceptual consequences of language cues on visual object recognition i.e., they focused only on the post-stimulus time interval. The mechanisms that could explain prestimulus effects of language on visual perception remain largely unknown.

Analysis of oscillatory activity provides an excellent opportunity to study language-driven prestimulus modulations in sensory areas. Based on previous human and animal studies, a candidate mechanism to carry sensory priors are low-frequency oscillations in the alpha/beta-band (Bastos et al. 2012; Michalareas et al., 2016; Arnal & Giraud, 2012; Bastos et al., 2015). Rhythmic brain activity in these frequency bands has been suggested to play a large variety of roles in top-down processing. For example, alpha synchrony has been associated with filtering of task irrelevant information and enhancement of neural representations during tasks involving attention, prediction, mental imagery and working memory (Mo et al., 2010; Mayer et al., 2015; Hari et al., 1997; Jensen et al. 2002). Similarly, beta oscillations have been implicated in perceptual expectations (Arnal & Giraud, 2012), online maintenance of cognitive states (Bressler & Richter, 2015; Engel and Fries, 2010) and (re)activation of task specific cortical networks (Spitzer & Haegens, 2017). Based on these findings, we hypothesized that any object recognition advantage for spoken words over natural sounds would be associated with a difference in cortical alpha/beta dynamics.

In the present study, we used a cue-picture matching task to test the hypothesis that language enhances visual object recognition by setting categorical priors via the modulation of alpha/beta oscillations. In contrast to previous studies, we (i) focused on the time interval preceding the onset of the visual object, targeting top-down signaling directly; and (ii) included words from participants’ first (L1) and second (L2) languages, in order to assess whether the previously reported label advantage extends to language systems acquired later in development

We hypothesized that, if the label advantage arises because words provide refined categorical priors to the visual system, then any differences in object recognition cued by words vs. natural sounds should be associated with modulations of oscillatory alpha/beta dynamics before the onset of the target picture. Importantly, we should also expect these oscillatory indices to be linked to behavioral performance.

## 2. Materials and methods

### 2.1. Participants

We tested a total of twenty-five Basque-Spanish bilingual speakers. Note that in earlier studies investigating the label advantage in object recognition, a sample size of 15 participants was sufficient to detect the behavioral label-advantage effect (Boutonnet and Lupyan, 2015). Participants were native speakers of Basque who began acquisition of Spanish after three years of age (13 females; age range 18-33, mean = 25.66, SD = 5.45, age of acquisition of Spanish = 4.23 y.o., SD = 1.33). All participants were right-handed, with no history of neurological disorders and had normal or corrected-to-normal vision. They received a payment of 10€ per hour for their participation. Before taking part in the experiment, all participants signed an informed consent form. The study was approved by the Basque Center on Cognition, Brain and Language (BCBL) Ethics Committee in compliance with the Declaration of Helsinki. Participants completed several language proficiency tests in both Spanish and Basque (see Table 1). First, participants were asked to self-rate their language comprehension (on a scale from 1 to 10, where 10 is a native-like level). All participants rated themselves as highly proficient in both Basque and Spanish. Participants also performed “LexTALE”, a lexical decision task (Izura et al., 2014; Lemhofer and Broersma, 2012) that tested their vocabulary knowledge. They obtained similarly high scores in both Spanish and Basque. In addition, participants had to name a series of pictures using vocabulary of increasing difficulty in both languages. Here as well, participants achieved native-range scores in both languages. Finally, all participants were interviewed by balanced bilingual linguists who rated them on a scale from 0 to 5: no participants had a score below 4 in either language.

**Table 1.** Measures of linguistic proficiency in Basque (L1) and Spanish (L2).

| Measure | Basque | Spanish |
| --- | --- | --- |
| Self-evaluation<br>(0-10) | 9.04 (0.16) | 9.39 (0.24) |
| LexTALE<br>Basque (0-50); Spanish (0-60) | 46.04 (2.67) | 54.09 (4.13) |
| Picture naming<br>(0-65) | 64.19 (1.47) | 63.38 (1.62) |
| Interview<br>(0-5) | 5 (0) | 4.95 (1.33) |

### 2.2. Stimuli

The visual stimuli comprised 50 pictures representing 10 animate (e.g., bird) or inanimate (e.g., camera) object categories. Each of these 10 categories was represented by 5 different highly recognizable images (.png extension, white background, 2000×2000 pixels): three color photographs obtained from online image collections, one normed color drawing (Rossion and Pourtois, 2004), and one “cartoon” image (Lupyan and Thompson-Schill, 2012). We selected different instances for each category in order to provide visual heterogeneity.

The audio stimuli comprised 10 words in Basque (L1), 10 words in Spanish (L2) and 10 natural sounds, each referring to one of the object categories. We recorded a Spanish-Basque female speaker producing both the Basque (L1) and Spanish (L2) words. Natural sound stimuli were downloaded from online libraries. Overall, the mean length of the audio stimuli was 0.8 ± 0.05 seconds (Word in L2, mean = 0.81 s, SD = 0.21; Word in L1, mean = 0.77 s, SD = 0.23; Natural Sounds, mean = 0.84 s, SD = 0.2).

In order to test that sounds and images were unequivocally identifiable, we asked a group of Basque-Spanish bilinguals (N=20), who did not take part in the main experiment, to view a selection of images and listen to a selection of sounds. They were told to name the visual and audio stimuli they perceived using the first noun that came to mind. For the present experiment, we only chose images and sounds which were identically named by all 20 participants. In total, experimental stimuli included 50 images from 10 categories, 10 words in Basque, 10 words in Spanish, and 10 natural sounds.

### 2.3. Procedure

The EEG experiment was run in a soundproof electrically shielded chamber with dim lighting. Participants sat on a chair, about sixty centimeters in front of the computer screen. Stimuli were delivered using PsychoPy software (Peirce, 2007). We followed the procedure illustrated by Boutonnet and Lupyan (2015). Participants completed a cued-picture recognition task composed of 300 trials (see Fig.1). On each trial, a fixation point appeared at the center of the screen for one second, then participants heard an auditory cue: either a word in L1, (e.g., *igela*, “frog”), a word in L2 (e.g., *rana*, “frog”) or a natural sound (e.g., a croak).

**Figure 1.**
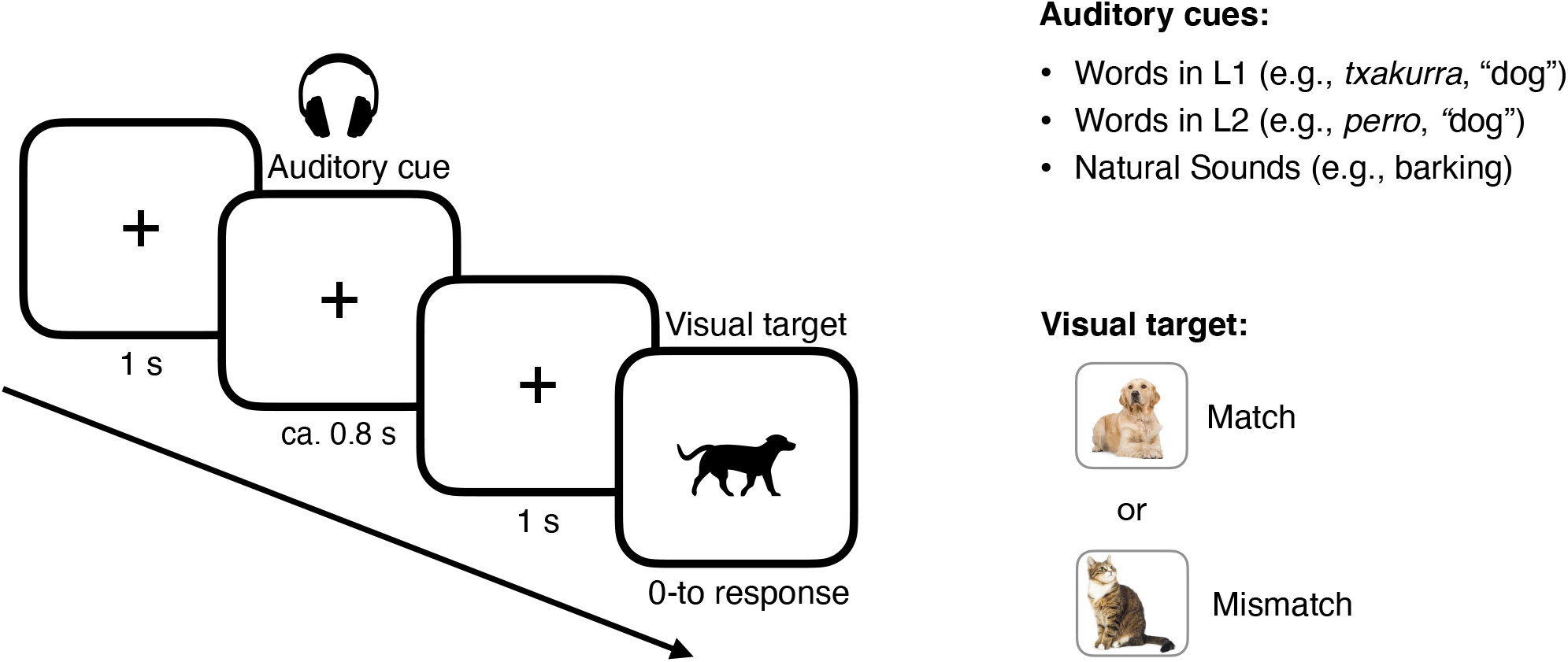
Cue-picture matching task. Participants were presented with auditory cues (words in L1, words in L2, natural sounds) and asked to evaluate whether the subsequent visual target did or did not match the auditory cue.

One second after cue offset, a picture appeared on the screen, and participants had to to indicate whether the picture did or did not match the auditory cue at the category level by pressing one of two buttons, “yes” or “no”, on the keyboard. The picture remained on screen until the participant responded. The picture matched the auditory cue in 50% of trials (congruent trials); in the other 50%, there was a mismatch (incongruent trials). In the case of incongruent trials, the picture that appeared on screen belonged to a different category. Stimuli presentation was randomized for each participant. The entire experiment lasted 40 minutes on average.

### 2.4. EEG recording

Electrophysiological activity was recorded from 27 electrodes (Fp1/2, F7/8, F3/4, FC5/6, FC1/2, T7/8, C3/4, CP1/2, CP5/6, P3/4, P7/8, O1/2, F/C/Pz) positioned in an elastic cap (Easycap) according to the extended 10–20 international system. All sites were referenced online to the left mastoid (A1). Additional external electrodes were placed on the right mastoid (A2) and around the eyes (VEOL, VEOR, HEOL, HEOR) to detect blinks and eye movements. Data were amplified (Brain Amp DC) with a filter bandwidth of 0.01-100 Hz, at a sampling rate of 250 Hz. The impedance of the scalp electrodes was kept below 5 kΩ; eye electrode impedance was kept below 10 kΩ.

### 2.5. EEG preprocessing

All EEG data analysis was performed using Matlab 2014 with the Fieldtrip toolbox (Oostenveld et al., 2011; http://www.fieldtriptoolbox.org) and R (R Core Team, 2015; http://www.r-project.org). For data visualization, we used Matlab or FieldTrip plotting functions, R and the RainCloud plots tool (Allen et al., 2019). The recordings were re-referenced off-line to the average activity of the two mastoids. Epochs of interest were selected based on cue type (words in L1, words in L2, natural sounds) and congruency (match, mismatch), resulting in six different sets of epochs, computed from −3 s to 1.5 s with respect to image onset.

Trials in which subjects provided incorrect responses in the behavioral task were removed from the analysis. Spatial-temporal components of the data containing eye and heart artifacts were identified using independent component analysis and subsequently removed. Overall, we removed an average of 2.14 components per subject. We then identified epochs containing additional ‘muscle’ and ‘eye blink’ artifacts using an automatic artifact detection procedure (z-value threshold = 12). Trials selected as possibly contaminated by artifacts were visually inspected and removed (~8%). Finally, we removed a few additional trials containing artifacts using a visual inspection procedure (~0.11%). Three participants were excluded from the analysis because more than 25 % of trials were rejected.

### 2.6. Statistical analysis

#### 2.6.1. Behavior

We used the R environment (version 4.0.0; R Core Team 2020) and lme4 package (Bates et al., 2014) to perform mixed effect regression on reaction time data, following a procedure similar to that illustrated in Boutonnet and Lupyan (2015). Predicted reaction times (calculated from the onset of the target image until the participant’s response) were computed by fitting the model with cue-type (words in L1, words in L2, natural sounds), congruency (match, mismatch), and their interaction as fixed factors, and by adding by-subject random slopes for the effect of cue type and congruency. Subsequent pairwise comparisons were performed using estimated marginal means (Bonferroni-corrected for multiple comparisons) with emmeans (Lenth, 2018). Because no reliable interaction was detected, post-hoc comparisons were based on a model with the same syntax as the one presented above but excluded the interaction term, in order to facilitate the interpretability of post-hoc analysis. Accuracy was not analyzed statistically because it was near ceiling (98%). For the analysis of behavioral data, we excluded the same three participants that were excluded from the EEG analysis. Moreover, we excluded all incorrect trials (1.88%), as well as a few trials in which participants’ responses exceeded 3 s (0.28%). These trials were also excluded from the EEG analysis. Reaction times were log-transformed to improve normality.

#### 2.6.2. Spectral power

A time-frequency analysis of artifact-free EEG trials was performed. Before applying spectral decomposition, the latency of each epoch was reduced to −1.5 s to 0.5 s with respect to image onset. The time-varying power spectrum of single trials was obtained using a Hann sliding window approach (0.5 s window, 0.05 s time steps) for the frequency range between 0 and 30 Hz, zero-padded to 1 s for a frequency resolution of 1 Hz. We focused on oscillatory activity up to 30 Hz because top-down processes are often associated with oscillations in this frequency band, while higher frequencies are linked to bottom-up processing (e.g., Bosman et al., 2012). For the statistical analysis, we computed a single power spectral density estimate for each participant, channel, frequency, and epoch by averaging the spectral estimates centered on the −0.75 s to −0.25 s time interval. We selected this time-interval to obtain more accurate spectral estimates, as activity here is largely uncontaminated by activity evoked by the preceding auditory event or subsequent visual stimulus.

#### 2.6.3. Grand-average power spectrum

In order to compute the power spectrum, we combined spectral estimates for congruent and incongruent trials for each cue-type condition, resulting in three different data sets (words in L1, words in L2, natural sounds). Note that time-frequency representations for congruent and incongruent conditions should be indistinguishable during the prestimulus time window since subjects had no way of anticipating the trial type. Spectral estimates were then averaged over trials, participants, channels, and cue-type conditions, resulting in a single value for each of the 30 frequency bins (i.e., the grand-average power spectrum). A peak-finding algorithm was used to identify spectral peaks as local maxima in the grand-averaged power spectrum. Two peaks, one at 10 Hz and one at 18 Hz emerged from this analysis (Fig. 2A). Based on these peaks, frequencies of interest (FOI) were obtained as the average of the frequency peaks ±1 Hz: that is, 9-11 Hz and 17-19 Hz respectively (Fig. 2B). We refer to these band estimates as alpha and beta band power, respectively. The topographical distribution indicates that these frequency peaks were larger over posterior electrodes for both the alpha (electrodes showing the greater effect: O1, O2, P8; mean = 9.91 μV2, SD = 1.88) and beta (electrodes showing the greater effect: O1, O2, P7; mean = 2.39 μV2, SD = 0.1) frequency bands (Fig. 2C).

**Figure 2.**
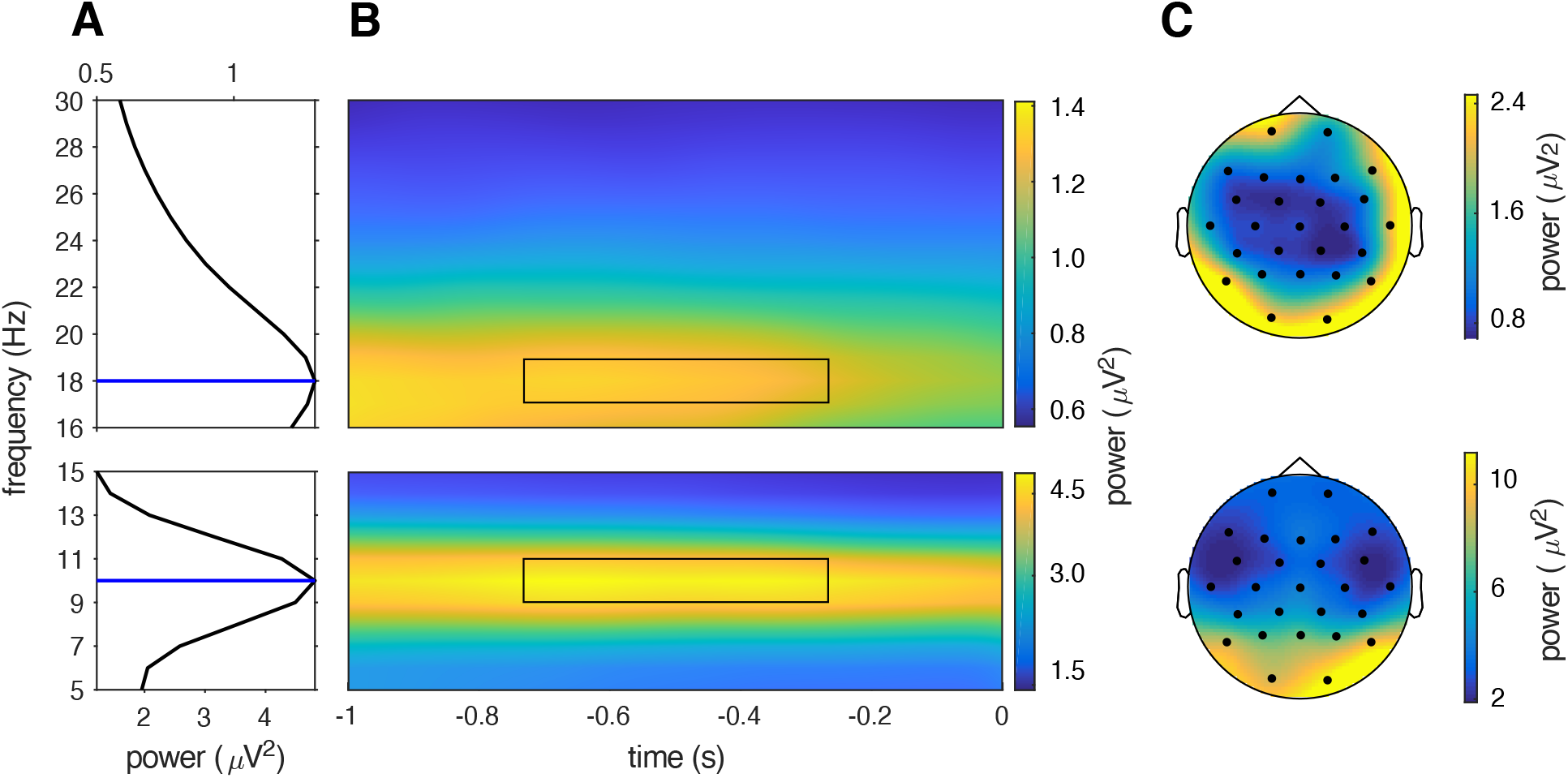
Time-frequency peaks and topographies. A) Alpha and beta peaks in the grand-average raw power spectrum of all epochs across conditions, during the −0.75 −0.25 pre-target time interval. The blues lines indicate the power peak as local maxima. B) Time-frequency representation of grand-averaged data for the alpha and beta-band, in the 1 s time window between the offset of the auditory cue and the onset of the image. The black rectangle denotes the timefrequency interval selected for the statistical analysis. C) Topography of the time-frequency interval of interest.

#### 2.6.4. Prestimulus spectral differences between cues

Spectral estimates for each cue-type (words in L1, words in L2, natural sounds) were averaged over trials. To reduce individual differences in overall EEG power, normalization was applied by converting the time-frequency power for each condition into percent signal change relative to the average power over all three conditions and channels, as performed by Bogaerts et al., (2020). This procedure removes individual differences in signal power, without distorting the relative magnitudes of the conditions, i.e., it functions as a baseline correction, when an appropriate baseline interval is not available. In order to test whether time-frequency representations in the prestimulus time-window differed across cue types, a non-parametric approach was selected (Maris and Oostenveld, 2007). For each FOI, we implemented a cluster-based permutation test based on a dependent sample F-test with the spectral data for each type of cue (words in L1, words in L2, natural sounds) as the dependent variable. This approach is equivalent to a one-way ANOVA but allowed us to account for the spatial correlation between electrodes (i.e., no a priori region of interest needs to be defined). The minimum number of neighboring electrodes required for a sample to be included in the clustering algorithm was set at 2. The cluster threshold F-value (or t-value) was set at an alpha value at the 85th percentile of their respective distributions. Note that this parameter does not impact the false alarm rate of the test. Rather, it sets a cluster threshold for determining when a sample should be considered as a candidate member of a cluster. Small cluster thresholds usually favor the detection of highly localized clusters with larger effect sizes, while larger cluster thresholds favor clusters with larger spatio-temporal extents but exhibit greater diffusion of the effect (Maris and Oostenveld, 2007). Because alpha and beta rhythms usually emerge at the network level, we selected a relatively large cluster threshold, i.e., capturing what appears to be a more globally distributed effect. The number of permutations for the randomization procedure was set at 100000. The critical alpha-level to control the false alarm rate was the standard α = 0.05. All resulting p-values were Bonferroni corrected for the number of FOIs. For each FOI, one significant cluster was detected. In order to assess the directionality of the effect, post-hoc non-parametric pairwise comparisons were applied. Specifically, power values for each cue-type condition were averaged over all electrodes belonging to the significant cluster and compared pairwise using paired t-tests. The alpha-level for the three post-hoc t-tests was Bonferroni corrected for the number of comparisons. This procedure was applied to each FOI separately.

For both the alpha and beta band, post-hoc t-tests revealed that brain data elicited by symbolic cues (words in L1 and L2) come from a similar probability distribution, while both significantly differed from brain activity elicited by natural sounds. This motivated us to pool the data from the spoken word conditions and contrast this average with that from the natural sound condition using a dependent sample cluster-based t-test (using the same parameters as for the test based on the F-statistic). This comparison gave one significant cluster for each FOI, respectively including 23 (alpha) and 21 (beta) out of 27 electrodes. The power over these clusters served for the analysis of brain-behavior correlations.

#### 2.6.5. Brain-behavior correlations

In order to investigate the link between prestimulus brain rhythms and behavior, correlation analyses were performed across participants. The correlation analyses reported below included only congruent trials, in which the auditory cue matched the object picture. The same analysis pipeline was applied for each FOI.

Trials were averaged, providing a single alpha, beta, and RT measure for each participant in each condition. To ensure the correlation was not driven by differences between conditions, participants’ values were z-scored within conditions. The three conditions were then averaged, providing an alpha-RT and beta-RT pair for each participant. Spearman correlations between these pairs were then computed for each FOI. To assess the statistical properties of the alpha and beta correlations, we bootstrapped the data over participants. We performed this 100000 times, generating a distribution of bootstrap values. Following Efron and Tibshirani (1986), we computed the percentile bootstrap 95% confidence intervals, and used this distribution to perform statistical tests to determine the difference between the observed correlation coefficients and zero. We finally conducted a two-sample bootstrap test to evaluate the difference between the alpha-RT and beta-RT correlation coefficients (Efron and Tibshirani, 1986).

## 3. Results

### 3.1. Effect of cues on visual object recognition

We first analyzed accuracy. Overall, accuracy was high (98%) and similarly distributed across the three conditions (words in L1 = 98%; words in L2 = 99%, natural sounds = 97%). Participants were clearly at ceiling, so we focused on the analysis of reaction times. Analysis of reaction time responses showed a main effect of Cue-Type (χ^2^(2) = 31.9500, p < 0.001) (Fig. 3). This was subsequently unpacked via post-hoc comparisons. Pairwise comparisons using estimated marginal means showed that object images preceded by symbolic cues in both L1 and L2 were identified faster than images preceded by natural sounds (words in L1 – natural sounds: Δ = −0.08, SE = 0.01, p < 0.001; natural sounds – words in L2: Δ = 0.06, SE = 0.01, p < 0.001). On the other hand, the pairwise effect between words in L1 and words in L2 did not reach the significance threshold (words in L1 – words in L2: Δ = −0.02, SE = 0.01, p = 0.06). As in previous studies, we also observed a main effect Congruency (χ^2^(1) = 7.0329, p < 0.01), with matching cue-picture pairs leading to faster responses than mismatching pairs. No reliable Cue-Type by Congruency interaction was detected (χ^2^(2) = 1.5310, p = 0.46).

**Figure 3.**
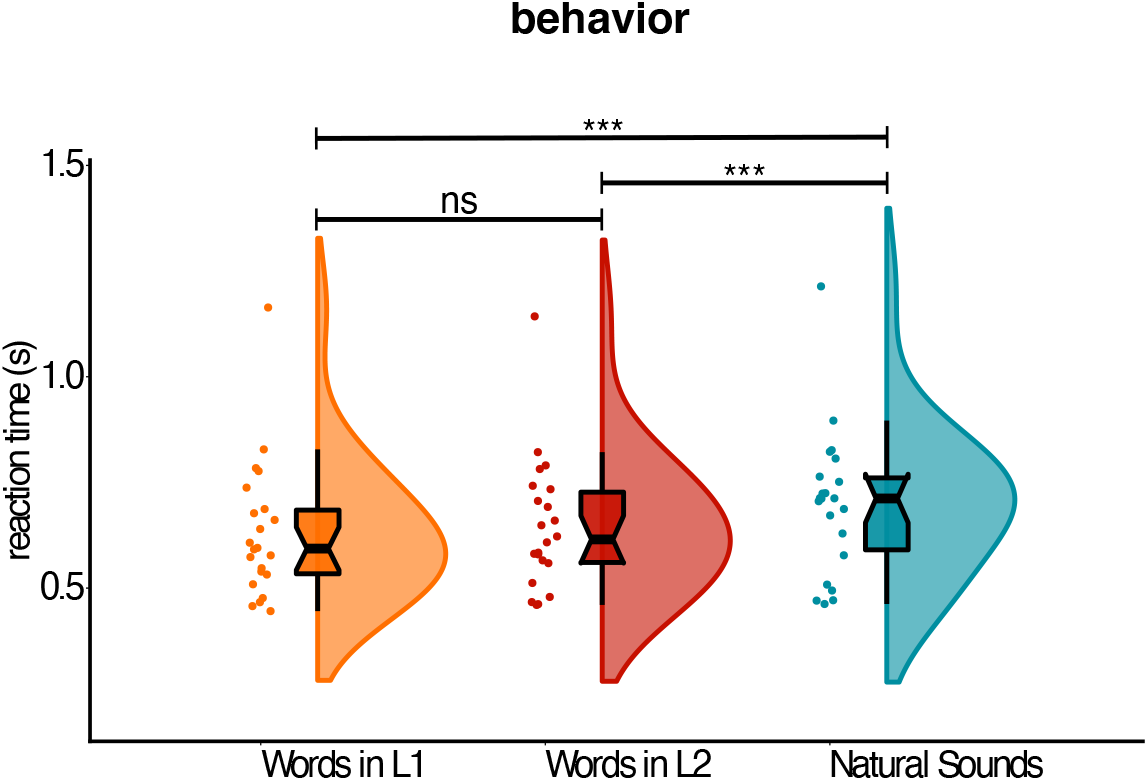
Behavioral results. Mean reaction times (correct trials only) showing the main effect of cue-type on visual object recognition performance. Raincloud plots show probability density. The center of the boxplot indicates the median, and the limits of the box define the interquartile range (IQR = middle 50% of the data). The notches indicate the 95% confidence interval around the median. Dots reflect individual subjects.

### 3.2. Effect of cues on prestimulus alpha rhythms

Differences between spectral power elicited by the three cue-type conditions were assessed using a cluster-based F-test for alpha and beta FOIs separately, focusing on the prestimulus interval. From the analysis of the alpha rhythm, one significant cluster was detected (p < 0.01, Bonferroni-corrected for the two FOIs) including several electrodes across the entire scalp (Fig. 4A). In order to assess the directionality of the effect, spectral power for each type of cue was averaged over all the electrodes belonging to the significant cluster and compared pairwise via t-tests. Pairwise comparisons showed that words in L1 and L2 both led to increased alpha power compared to natural sounds (t(21) = 4.57, p < 0.001 Bonferroni-corrected; t(21) = 5.48, p < 0.001 Bonferroni-corrected, respectively) (Fig. 4A). No significant difference was detected between words in L1 and L2 (t(21) = −1.70, p = 1 Bonferroni-corrected).

**Figure 4.**
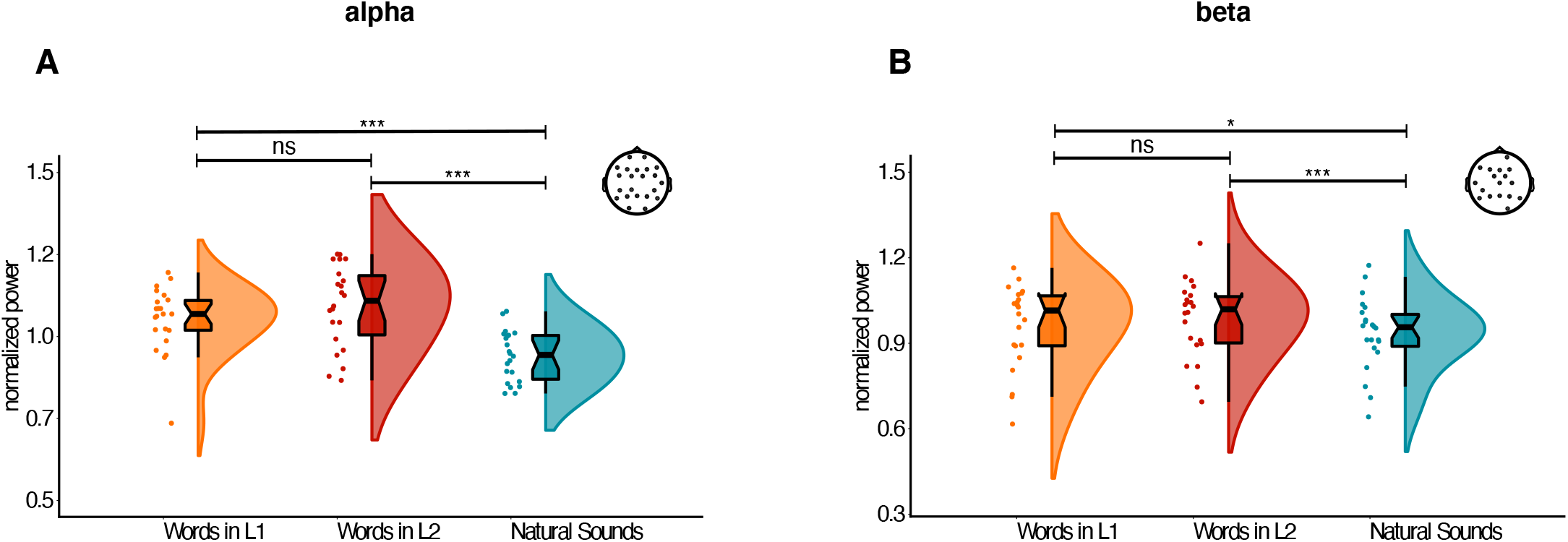
EEG results. Effect of cues on pre-target alpha (A) and beta power (B) averaged over the electrodes belonging to the significant cluster. Conventions for the plot are the same as in Figure 2. The electrodes belonging to the cluster are illustrated at the top-right of each figure.

### 3.3. Effect of cues on prestimulus beta rhythms

Beta band analysis revealed a pattern of results similar to the alpha rhythm analysis. The cluster-based F-test detected one cluster (p < 0.01, Bonferroni-corrected for number of FOIs) (Fig. 4B). Spectral power in the beta frequency range was averaged over the electrodes of the significant cluster for each type of cue separately and compared pairwise via t-tests. Beta power was larger when images were preceded by words in L1 and L2 compared to natural sounds (t(21) = 2.68, p = 0.04 Bonferroni-corrected; t(21) = 4.68, p < 0.001 Bonferroni-corrected, respectively), while no significant difference emerged when comparing words in L1 and L2 (t(21) = −1.67, p = 0.33 Bonferroni-corrected) (see Fig. 3B).

### 3.4. Relation between prestimulus alpha/beta rhythms and visual object recognition

The results reported so far point to a possible role for alpha and beta rhythms in supporting the label-advantage in object recognition. We further explored the relation between prestimulus alpha/beta oscillations and visual object recognition by correlating prestimulus spectral power and reaction times across participants.

Individual estimates for power and reaction times were correlated using Spearman rank correlations. This method was selected because reaction times and beta power data significantly deviated from normality, as shown by a Shapiro-Wilk normality test (reaction time: W = 0.88, p = 0.01; alpha power: W = 0.92, p = 0.09; beta power: W = 0.89, p = 0.02). We observed that both prestimulus alpha and beta power correlated with reaction times in the object recognition task (Fig. 5A). Yet, these effects exhibited opposite directionality: alpha estimates were negatively correlated (Spearman’s rho = −0.34, p = 0.13), while beta power estimates were positively correlated (Spearman’s rho = 0.33, p = 0.13). Though these correlations were not significant, the bootstrap percentile confidence intervals (Fig. 5B) provided marginal evidence suggesting that the alpha correlation was less than zero, as shown by the 95% confidence interval only minimally exceeding zero (zero lies at the 93rd-percentile of the alpha bootstrap distribution). This can be expressed as a p-value using the bootstrap distribution to test the one-tailed hypothesis that the alpha-RT correlation is less than 0 (p = 0.067), again providing marginal evidence that the alpha-RT correlation was negative. Similarly, the bootstrap analysis provided evidence supporting a positive beta-RT relationship, with zero appearing at the 4.9th-percentile, close to the border of the 95% confidence interval (Fig. 5B) and a significant difference from zero using the one-tailed hypothesis test (p = 0.049). Building on this evidence for opposing effects of alpha and beta power on RT, we explicitly tested whether these correlations were in fact different. Figure 5B shows that the observed rho values for the alpha-RT and beta-RT correlations fall outside each other’s 95% confidence intervals. A two-tailed two sample bootstrap test was applied, which revealed that indeed the alpha-RT and beta-RT correlations are significantly different (p = 0.012), thus supporting the interpretation that these oscillations exert different effects on behavior.

**Figure 5.**
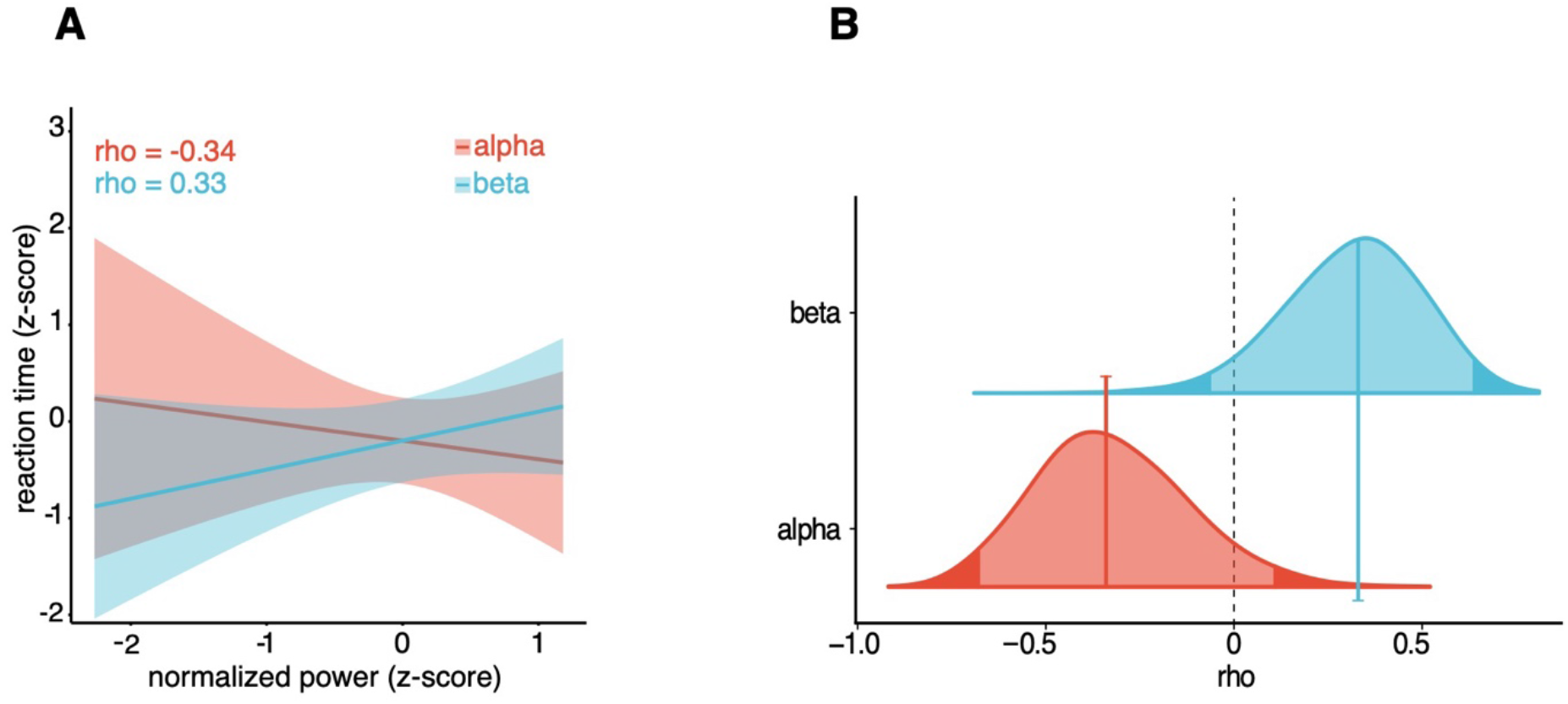
Correlation analyses. A) Correlation between alpha/beta power and reaction time (RT). Error bars represent 95% confidence interval. B) Bootstrap distributions of alpha/beta Spearman rho values.

## 4. Discussion

Several studies have reported that spoken words can boost visual recognition of object categories, but the neural mechanisms underlying such facilitation is not well established. It has been suggested that effects of language on visual perception arise at early stages of sensory processing; specifically, via the amplification of category-specific priors in sensory regions. In the present study, we investigated the prestimulus effect of language on visual perception, testing the hypothesis that neural oscillations can serve as mechanisms to carry language-generated priors about incoming object categories.

To test this hypothesis, we used EEG to measure prestimulus brain activity and characterize the oscillatory dynamics underlying the label-advantage in object recognition. We reasoned that, if objects are recognized faster because spoken words provide more refined categorical priors than natural sounds, then these cues should differentially modulate prestimulus oscillatory activity in the alpha/beta bands; and this oscillatory index should be linked to object recognition performance.

We first replicated the previously reported label-advantage and showed that this behavioral effect persisted even when words were presented in a second language. This suggests that verbal symbols deploy more accurate knowledge representations than natural sounds against which incoming inputs can be compared. Importantly, the reported behavioral advantage for spoken words was associated with an increase in the power of alpha and beta rhythms in the time interval between the offset of the cue and the onset of the target object. Such synchronization points to a possible functional role for alpha and beta neural rhythms in the label advantage in object recognition.

While the wide scalp distribution of these oscillatory indices might point to global alpha/beta network states, we found more pronounced peaks of activity over posterior electrodes. This was particularly evident in the alpha frequency-band (Fig. 2C) and suggests alpha waves may reflect local oscillatory states originating in occipital regions, in line with findings from the monkey literature (e.g., Bollimunta et al., 2008; Mo et al., 2011a). Enhancement of alpha oscillations in occipital regions have been largely reported when top-down knowledge is directed by a cue towards a specific feature or direction (Worden et al., 2000; Snyder and Foxe, 2010). At least two non-mutually exclusive theoretical accounts have been advanced to explain this effect. Some recent proposals posit that enhancement of neural alpha synchronization in task-relevant regions leads to excitatory effects reflecting selective amplification of neural representations of object categories (Palva and Palva, 2007; Klimesch, 2012; van Kerkoerle et al., 2014; Mo et al., 2011a). For instance, M/EEG studies have reported that alpha power increases in grapheme-processing regions with the predictability of letter identity (Mayer el al., 2016); and in the posterior cortex when meaningful hints precede the discrimination of ambiguous images (Samaha et al., 2018). Similarly, biophysical models indicate that enhancement of prestimulus alpha waves can improve detection performance by increasing excitation of pyramidal cells, rendering the network state less stable and thus facilitating the activation of a stimulated assembly (Lundqvist et al. 2013). In line with these studies, we suggest that alpha oscillations can serve as a mechanism to carry language-generated priors about the structure of visual objects. This interpretation is also partially supported by the negative correlation between individual alpha power and reaction times, showing that participants with higher prestimulus alpha power were overall faster at recognizing visual objects.

Another prominent view is that enhanced alpha power reflects states of inhibition and filtering of task irrelevant information (Jensen and Mazaheri, 2010; Klimesch et al., 2007). For instance, when attention is directed towards a target in one side of space, posterior alpha-band power increases at electrodes over the hemisphere ipsilateral to the target (Worden et al., 2000; Thut et al., 2006). According to this view, increased alpha oscillations reflect suppression of cortical areas not involved in the task. The alpha effect in our study was right-lateralized and might reflect inhibition of right-posterior regions to gate sensory information processing to the left-posterior network, where language-perception interactions usually take place (Gilbert et al., 2008; Mo et al., 2011b). However, this interpretation would also predict alpha desynchronization over left-posterior regions to increase excitability and enhance stimulus processing (Jones et al., 2000; Klimesch et al., 2007). Since we did not find any evidence for the latter effect, we consider unlikely that alpha synchronization acted as an inhibitory filter in the current study.

A novel result of our study in contrast with similar earlier studies was the differential beta-band modulations that resulted from spoken word vs. natural sound cues. Recent proposals suggest that beta oscillatory activity reflects endogenously driven transitions from latent to active cortical representations of objects categories (Spitzer and Haegens, 2017), as well as the binding of neurocognitive network elements associated with a given neural representation (Bressler and Richter, 2015). Under these accounts, beta synchronization provides “a flexible scaffolding that sets up functional neuronal ensembles through temporary synchronization of content-coding cell populations” (Spitzer and Haegens, 2017). We speculate that the difference in beta modulations for spoken words vs. natural sounds may reflect a difference in the content of the (re)activated conceptual states – and more importantly, in the amount of retrieved conceptual dimensions, e.g., the size of the neurocognitive network state (Bressler and Tognoli, 2006). Behavioral and eye tracking experiments have indeed shown that spoken words activate a rich network of features during lexical processing (e.g., Huettig and Altmann, 2005). As a consequence, processing words might lead to the retrieval of knowledge dimensions that go beyond the purely sensory features of objects, such as conceptual, grammatical and lexical information. This is partially in line with human and monkey studies showing that beta synchronization carries supramodal information about object categories (Antzoulatos and Miller, 2014, 2016; Wutz et al., 2018).

The correlations we found between the individual magnitude of prestimulus alpha/beta power and behavioral performance indicate that object recognition improved when prestimulus alpha power increased but slowed down with increases in beta synchronization. Although these correlations did not reach the threshold of significance, possibly due to the relatively small sample size of the current study, follow-up analyses provided some evidence that the alpha-RT correlation was negative, while the beta-RT relationship was positive. This result suggests that neural synchronization in these frequency bands might not be incidental to object recognition performance. Interestingly, bootstrap analyses also revealed that the alpha-RT and beta-RT correlations were different. This finding points to a possible division of labor between alpha and beta rhythms in top-down signaling during language-mediated visual object recognition, where alpha rhythms might function to amplify neural representations of visual object categories, while beta-frequency synchronization may (re)activate or maintain the neurocognitive network states elicited by the auditory cue. However, given that these correlations did not reach significance, any interpretation of these results remains speculative and awaits further investigation.

The present findings inform the broad debate on whether language shapes perception at early or late stages of perceptual processing. Evidence for the former account comes primarily from EEG studies showing that language affects visual processes by modulating early ERP components such as the P1 (Boutonnet and Lupyan 2015; Noorman et al., 2018) and N170 (Landau et al., 2010). However, studies focusing on post-stimulus activity are also coherent with a later semantic or decisionmaking account. Indeed, post-stimulus differences, even if very early, could still emerge from rapid feed-forward integration of visual and linguistic information (Thierry et al., 2009). The prestimulus modulations of alpha waves in posterior regions reported here provide complementary support for the idea that linguistic influences on perception arise at early stages of sensory processing and suggest that the mechanism underling such influences is the amplification of sensory priors in posterior regions via alpha-band synchronization.

Finally, our study included a novel manipulation not considered in previous studies on categorization: the inclusion of L2 words as auditory cues. Our participants were highly proficient Basque-Spanish bilinguals, with comparable levels of proficiency in both languages, who had acquired their L2 later in development. The effect of top-down processing in bilinguals is currently debated, and largely dependent on factors like proficiency (Kaan, 2014; Hopp, 2013) and age of acquisition (Molinaro et al., 2017). Although it is commonly believed that bilinguals access a semantic system common to both languages (e.g., Caramazza and Brones, 1980), recent studies have suggested that top-down processing may be reduced in a second language because of reduced access to perceptual memory resources (e.g., Hayakawa and Keysar, 2018), which are known to play an important role in the generation of visual expectations (Hindy et al., 2016). We found comparable behavioral and neural responses after L1 and L2 words cuing visual object recognition. This result is in line with the idea that both languages access common conceptual representations and provide similar types of top-down guidance to the visual system.

However, our results show that L1 and L2 words both affect visual processing differently than natural sounds, challenging the hypothesis that such cues provide similar top-down priors to visual regions. It is worthwhile considering why a word cue results in better visual object recognition than a natural sound. It has been proposed that symbols are extremely effective in compressing semantic information in a format that transcends within-category differences, thus leading to the amplification of those features which are relevant for distinguishing between exemplars of different categories. By contrast, natural sounds are inevitably linked to specific sources (e.g., the barking of a dog may trigger the representation of a specific exemplar of a dog), thus being less effective at cuing categorical states (Edmiston and Lupyan, 2015). Interestingly, ascribing labels to experiences has also been shown to enhance other cognitive functions, such as the retention of items in visual working memory (Souza and Skóra, 2017), learning novel categories (Lupyan et al., 2007), and perceptual categorization across sensory modalities (e.g., Miller et al., 2018). These findings indicate that language acts as a powerful tool for compressing information, facilitating different operations important to a multitude of human cognitive processes (Clark 2012). Future studies should investigate whether similar oscillatory mechanisms are employed to support these language-augmented cognitive functions.

## Acknowlegments

This research was supported by the Spanish Ministry of Economy and Competitiveness (MINECO), Agencia Estatal de Investigacion (AEI), and Fondo Europeo de Desarrollo Regional (FEDER) (PSI2015–65694-P and RTI2018-096311-B-I00 to NM) and by the Basque government (grant PI_2016_1_0014 to NM). The authors acknowledge financial support from the Spanish Ministry of Economy and Competitiveness, through the “Severo Ochoa” Programme for Centres/Units of Excellence in R&D (SEV-2015-490). Work by Piermatteo Morucci received support from “la Caixa” Foundation (ID 100010434) through the fellowship LCF/BQ/IN17/11620019, and the European Union’s Horizon 2020 research and innovation programme under the Marie Skłodowska-Curie grant agreement no. 713673. We wish to express our gratitude to the BCBL lab staff and the research assistants who helped to recruit the participants and collect the data. We also thanks to Magda Altman and Jose Pérez-Navarro for their helpful comments on the manuscript.

## Conflict of Interests

The authors declare no competing financial interests.

## Data statement

The behavioral and EEG data reported in this paper, experimental materials, and analysis code are available via the Open Science Framework (https://osf.io/ntg9q/). This study has not been preregistered.

## Authors contribution

**Piermatteo Morucci**: Software, Validation, Formal analysis, Resources, Data Curation, Visualization, Project administration, Writing–original draft, Writing–review & editing. **Francesco Giannelli**: Conceptualization, Methodology, Software, Investigation, Resources, Data Curation, Project administration, Writing–review & editing. **Craig Richter**: Formal analysis, Supervision, Writing–review & editing. **Nicola Molinaro**: Supervision, Funding acquisition, Project administration, Writing–review and editing.

## Notes

### Competing Interest Statement

The authors have declared no competing interest.

